# GSMN-ML- a genome scale metabolic network reconstruction of the obligate human pathogen *Mycobacterium leprae*

**DOI:** 10.1101/819508

**Authors:** Khushboo Borah, Jacque-Lucca Kearney, Ruma Banerjee, Pankaj Vats, Huihai Wu, Sonal Dahale, Manjari K Sunitha, Rajendra Joshi, Bhushan Bonde, Olabisi Ojo, Ramanuj Lahiri, Diana L. Williams, Johnjoe McFadden

## Abstract

Leprosy, caused by *Mycobacterium leprae*, has plagued humanity for thousands of years and continues to cause morbidity, disability and stigmatization in two to three million people today. Although effective treatment is available, the disease incidence has remained approximately constant for decades so new approaches, such as vaccine or new drugs, are urgently needed for control. Research is however hampered by the pathogen’s obligate intracellular lifestyle and the fact that it has never been grown *in vitro*. Consequently, despite the availability of its complete genome sequence, fundamental questions regarding the biology of the pathogen, such as its metabolism, remain largely unexplored. In order to explore the metabolism of the leprosy bacillus with a long-term aim of developing a medium to grow the pathogen *in vitro*, we reconstructed an *in silico* genome scale metabolic model of the bacillus, GSMN-ML. The model was used to explore the growth and biomass production capabilities of the pathogen with a range of nutrient sources, such as amino acids, glucose, glycerol and metabolic intermediates. We also used the model to analyze RNA-seq data from *M. leprae* grown in mouse foot pads, and performed Differential Producibility Analysis (DPA) to identify metabolic pathways that appear to be active during intracellular growth of the pathogen, which included pathways for central carbon metabolism, co-factor, lipids, amino acids, nucleotides and cell wall synthesis. The GSMN-ML model is thereby a useful *in silico* tool that can be used to explore the metabolism of the leprosy bacillus, analyze functional genomic experimental data, generate predictions of nutrients required for growth of the bacillus *in vitro* and identify novel drug targets.

**Author Summary:** *Mycobacterium leprae*, the obligate human pathogen is uncultivable in axenic growth medium, and this hinders research on this pathogen, and the pathogenesis of leprosy. The development of novel therapeutics relies on the understanding of growth, survival and metabolism of this bacterium in the host, the knowledge of which is currently very limited. Here we reconstructed a metabolic network of *M. leprae*-GSMN-ML, a powerful *in silico* tool to study growth and metabolism of the leprosy bacillus. We demonstrate the application of GSMN-ML to identify the metabolic pathways, and metabolite classes that *M. leprae* utilizes during intracellular growth.

## Introduction

The mycobacterial genus includes two of the greatest scourges of humanity: *Mycobacterium tuberculosis* and *M. leprae*, responsible for tuberculosis (TB) and leprosy respectively. Leprosy is one of the oldest plagues of mankind yet remains prevalent in developing countries. In 1981, the World Health Organization (WHO) recommended multidrug treatment (MDT) of disease with dapsone, rifampicin and clofazimine. Implementation of MDT achieved a 98% cure rate leading to reduction of prevalence by a remarkable 45% within three years. Yet, although MDT reduced global prevalence of leprosy, annual new case detection has remained fairly constant over the last decade [1]. Problems with control include the extremely lengthy drug treatment complicated by reactions that can lead to nerve damage and worsening of symptoms. Relapse of infection can also occur after treatment if the MDT therapies fail to clear drug resistant bacteria [2]. Additionally, new cases occurring in children, reported at 9.2% in 2013, have not reduced, indicating that active transmission continues [1], [2]. According to the WHO reports, 210,758 new cases of leprosy were detected in 136 countries in 2015 with the highest number of cases recorded in the developing nations of South-east Asia [3]. New approaches are clearly needed for case-finding, diagnosis, treatment and tackling social stigma of this devastating disease [3], [4].

One of the most striking facts about leprosy is how different it is from the disease caused by its related pathogen, *M. tuberculosis*. Both *M. tuberculosis* and *M. leprae* are intracellular pathogens that replicate primarily in macrophages but whereas the TB bacillus appears to primarily infect macrophages (and possibly also dendritic cells), the leprosy bacillus replicates in many cell types but particularly Schwann cells and endothelial cells [5, 6]. Possibly as a consequence of their differing cell tropism, tuberculosis is predominantly a pulmonary inflammatory infection (although extra-pulmonary infections are common) characterized by strong cellular immune reactions; whereas leprosy causes both disseminated infections and localized infections [7], [8]. The replicative capabilities of the two pathogens are also different, with *M. tuberculosis* being able to grow both intracellularly and extracellularly; whereas *M. leprae* is an obligate intracellular pathogen. Additionally, *M. tuberculosis* is able to grow *in vitro*, in minimal medium containing a single carbon source such as glycerol, a nitrogen source, such as ammonium ions and a few essential mineral ions [9], whereas *M. leprae* has never been grown in axenic media, even with the addition of various growth supplements [10]. This latter feature severely hampers most biological studies of the pathogen. The replication rate of the pathogens also differs markedly. Although both are very slow growing organisms, *M. tuberculosis* replicates with a doubling time of about a day in optimal conditions whereas the doubling time of *M. leprae* in infected mouse is 10-14 days [11], [12]. The reasons for these differences in growth, virulence characteristics and cell tropism still remain unknown.

A major advances in the understanding the pathogenesis of both TB and leprosy was achieved with the publication of their complete genome sequences, first with *M. tuberculosis* in 1998 [13] and then *M. leprae* in 2001 [10]. Analysis of the *M. leprae* genome sequence revealed that the pathogen had undergone massive gene decay with only 1604 predicted open reading frames compared to 3959 in *M. tuberculosis*. The reduced genome is a consequence of multiple mutations in coding sequences leading to gene decay, and the accumulation of 1116 pseudogenes and thereby loss of their associated functions [12], [14]. The loss of so many genes raised the obvious possibility that the inability of *M. leprae* to grow in axenic culture is due to mutational loss of biosynthetic function and consequent reliance on importing complex nutrients from the host which are not available in artificial media, as has been found for several other intracellular parasites. However, analysis of the genome sequence of *M. leprae* demonstrated that the most of the anabolic capability of the pathogen, relative to *M. tuberculosis*, seems to be intact. The pathogen appears to have retained complete enzyme systems for synthesis of all amino acids, except for methionine and lysine [10]. The biosynthesis of purines, pyrimidines, nucleosides, nucleotides, vitamins and cofactors is also impaired [10]. Gene deletion and decay does however appear to have eliminated many important catabolic systems and respiratory systems. For example, as previously reported Cole *et al*., 2001, the aerobic respiratory chain of *M. leprae* is truncated due to loss of several genes in the NADH oxidase operon, nuoA-N, potentially eliminating the ability of the pathogen to produce ATP from the oxidation of NADH. Yet central carbon metabolism appears to be intact in the pathogen with an intact glycolytic pathway, pentose phosphate pathway (PPP), gluconeogenesis and the tricarboxylic acid (TCA) cycle. Thus, despite the availability of the complete genome sequence of *M. leprae*, the inability of the pathogen to grow in axenic culture remains a mystery. Also, the substrates utilized during growth of *M. leprae in vivo*, remain unclear. Recent research has also revealed intriguing capabilities of the leprosy bacillus to manipulate its host cell including its ability to dedifferentiate and reprogram its host Schwann cell to a stem-cell-like state in which the pathogen can proliferate more efficiently and which likely promotes its dissemination through the body [15].

The availability of full genome sequences does however allow reconstruction of genome-scale metabolic reaction networks for micro-organisms. These not only instantiate current knowledge of the metabolism of particular organisms; but may be used to analyze and integrate data, particularly functional genomic data, to generate novel predictions that may be subjected to experimental validation. Genome-scale metabolic networks have been constructed for many microbes including *M. tuberculosis* and *Mycobacterium bovis* where they have been used to identify novel metabolic pathways and examine their growth and likely substrate utilization within their host cells [16], [17], [18], [19]. In this study we constructed the first genome-scale network of *M. leprae*, GSMN-ML. We used the network to examine the organism’s catabolic and anabolic capabilities to gain insight into why the pathogen is unable to grow *in vitro*. We also used GSMN-ML to interrogate a RNA-seq dataset obtained from *ex vivo* grown *M. leprae* to probe the pathogen’s metabolic response to its host environment and analyzed the data using differential producibility analysis (DPA), a bioinformatic tool that translates transcriptomic data to predictions of metabolic responses [20].

## Materials and methods

### Ethics statement

All animal studies were performed under the scientific protocol number A-105 reviewed and approved by the National Hansen’s Disease Programs (NHDP) Institutional Animal Care and Use Committee, and were conducted in strict accordance with all state and federal laws in adherence with PHS policy and as outlined in The Guide to Care and Use of Laboratory Animals, Eighth Edition. Mice were euthanized by CO_2_ asphyxiation followed by cervical dislocation.

### *M. leprae* culture and growth conditions

Athymic nude mice (Envigo) were infected by inoculating 3 × 10^7^ *M. leprae*, strain Thai-53, in each hind footpad. At 4-5 months post inoculation mice were euthanized and footpad tissues removed. These tissues were either fixed in 70% ethanol for at least 48 h or minced and homogenized immediately for purification of viable *M. leprae* [21]: *in vivo M. leprae*. Viable bacteria were then also added to axenic medium 7H9 containing Caesitone (0.1% w/v), BSA (0.5% w/v), dextrose (0.75% w/v) and ampicillin (50µg/ml), and incubated at 33°C, 5% O_2_ for 48h. Following incubation, bacteria were pelleted and fixed in 70% ethanol: *in vitro M. leprae*. After fixation for at least 48h, ethanol was removed and *M. leprae* were treated with 0.1N NaOH to remove mouse tissues. Bacteria were adjusted to 2 × 10^9^/ml and 1 ml aliquots were pelleted and resuspended in 1 ml Trizol and stored at -20°>C. *M. leprae* in ethanol-fixed tissues were also purified, treated with NaOH, resuspended in Trizol and stored at -20°>C.

### RNA Purification and RNASeq Analysis

Bacterial lysis and RNA purification was performed using FastPrep-24 vertical homogenizer (MP BioMedicals), and FastPrep Lysing Matrix B tubes. After DNase treatment, an aliquot of sample RNA was converted to cDNA and analyzed for the presence of *M. leprae esxA* expression and DNA contamination [21]. Ribosomal RNA was depleted from total RNA preparations using Ribo-Zero rRNA Removal Kit according to manufacturer’s recommendations, and RNA quantity and quality was determined using NanoDrop 2000 and Agilent 2100 BioAnalyzer. Sequencing samples were prepared using the SOLiD® Total RNA-Seq Kit (Life Technologies, PN4445374) using 100 ng ribosome RNA-depleted and barcoded t-RNA (SOLiD™ RNA Barcoding Kit, Module 1-16 (Life Technologies, PN 4427046) according to the manufacturer’s protocol. Emulsion PCR and SOLiD sequencing, 75 base pairs single direction, were performed for the SOLiD™ 5500 System (Life Technologies, Thermo Fisher Scientific). Sequencing analysis was done using Bacterial RNA-seq Analysis Kit 1.0 on Maverix Analytic Platform (Maverix Biomics Inc.). Raw sequencing reads from SOLiD sequencing platform that were converted into FASTQ file format were quality checked for potential sequencing issues and contaminants using FastQC (FastQC). Adapter sequences, primers, Ns, and reads with quality score below 13 were trimmed using fastq-mcf of ea-utils [22] and Trimmomatic [23]. Reads with a remaining length of < 20 bp after trimming were discarded. Pre-processed reads were mapped to the *Mycobacterium leprae* TN genome (RefSeq Accession Number: NC_002677) using EDGE-pro [24]. Read coverage on forward and reverse strands for genome browser [25] visualization was computed using BEDtools [26], SAMtools [27] and UCSC Genome Browser utilities [23]. Read counts for RefSeq genes generated by EDGE-pro [25] were normalized across all samples and then used for differential expression analysis using DEseq [28]. Significant differentially expressed genes were determined by adjusted P-value with a threshold of 0.05. Fold change (log_2_) between samples were hierarchically clustered using Pearson correlation.

### Construction of GSMN-ML

The updated *M. tuberculosis* genome scale metabolic network (GSMN-TB_2) [16] created for H37Rv strain (supplementary File S1) was the basis of reconstruction of *M. leprae* metabolic model GSMN-ML (Supplementary File S3). The reconstruction of the network followed the methods as described previously [9], [16]. *M. leprae* (RefSeq ID: NC_002677) functional genes were identified using the online GenBank tool [29], and compared to the same list used for the development of the GSMN-TB of *M. tuberculosis* using the bi-directional best hit criteria to search for orthologs [30]. Using *M. tuberculosis* H37Rv (RefSeq ID: NC_000962) as a template, the metabolic network was reconstructed using Pathway Tools (Version 24.0) [31]. Non-functional pathways in, and gene essentiality data for *M. leprae* were identified and the model refined to remove any non-homologous pathways, identified by literature search and metabolic pathway data from previous mutagenesis study [32]. Surrey-FBA was used in the study for the flux balance analysis to further refine the model by iteration [33]. Single genes, and the respective pathways were removed and the model’s feasibility re-assessed. Only edits which did not render the model infeasible were used in the final product. If found to be essential, transport reactions were introduced for their product(s), so that the inactivation of these genes will be compensated by uptake of the predicted metabolites from the artificial media (presumed to be obtained from host in vivo). The full list of reactions, for GSMN-ML and GSMN-TB_2 models can be found in supplementary files S1 and S3 respectively.

### Flux balance analysis (FBA)

FBA simulations in this study were also performed with the SurreyFBA software as described previously [9], [33]. Simulations were conducted under steady-state conditions, using biomass production as the objective function to obtain the steady state theoretical fluxes through the reactions in the network that maximised biomass production. For FBA, substrate inputs for carbon and nitrogen sources were specified in the test medium that was used to constrain the simulations. External metabolites, except in the cases of those identified as being rate-limiting, were considered to be freely available and disposable. Rate-limiting external nutrients are bound by upper and lower flux rate limits, as described in the model’s set-up. FVA analysis were performed as described in Gevorgyan et al., 2011 to obtain the upper and lower flux limits through a particular reaction [33]. Gene essentiality predictions were tested using SurreyFBA software. The maximal theoretical growth rate of each in silico gene knock out was calculated by removing single genes from the network and performing FBA linear programming as described previously [9], [16], [33].

### Differential producibility analysis (DPA)

DPA seeks to translate transcriptomic signatures into metabolic fluxes using FBA to identify metabolites or metabolic pathways most affected by the changes in gene transcription and is described in detail by Bonde et al., 2011 [20]. RNA-seq data from *M. leprae*, strain Thai-53 cultivated in mice foot pads were used for DPA. DPA utilized a FBA-based metabolite expression analysis using glucose or glycerol as the carbon source input to GSMN-ML to calculate the maximal theoretical flux towards each metabolite in the network. For every metabolite, reactions involved were then ranked based on the expression values, with those ≥ 2 being highly expressed and those < 2 lowly expressed. A median value was assessed, creating a matrix of values for each metabolite. The same process is repeated with down-regulated reactions and the result is two lists of the most differentially produced metabolites for a given experimental environment. Statistical significance of the DPA is assessed by the following:

≥*0*, o*r*

≤*0, th* =1

*oth* =0

*E*=*S*/*N*

where RUm = median expected value of up-regulated metabolite

DUm = median expected value of down-regulated metabolite

O = observed value in real dataset

S = arbitrary score value

N = number of datasets

E = E-value

A significance threshold of E <0.15 was decided for further analysis

## Results

### Construction of GSMN-ML genome scale metabolic network

Our approach was to start with our previously-constructed metabolic model of *M. tuberculosis*, GSMN-TB [16] and step-by-step, replace all *M. tuberculosis* genes with orthologues from *M. leprae*. We used an updated version of GSMN-TB_2, which includes reactions for cholesterol catabolism (Supplementary File S1). We adjusted the biomass composition to reflect what is known about the macromolecular composition of *M. leprae*, for example, removing *M. tuberculosis*-specific components and adding the phenolic glycolipid (PGL1) which is a major antigen in leprosy. This entailed adding additional reactions for PGL1 synthesis. Also, *M. leprae* lacks N-glycosylated muramic acid in its peptidoglycan; and glycine is substituted in place of L-alanine [34], so this was reflected in the *in silico* peptidoglycan composition. Mannosyl β-1 phosphodolichol (MPD) of *M. tuberculosis* is a polyketide known to invoke strong T-cell response and pks12 [35], which is required for formation of MPD is a pseudogene in *M. leprae* (ML1437c) and therefore, the corresponding biosynthesis reactions were removed from the model. A small number of additional metabolic reactions that are found in *M. leprae* but absent in *M. tuberculosis*, were added to the network. These included ML2177, a putative uridine phosphorylase involved in nucleotide salvage, highlighting the previously described difference between nucleotide salvage in *M. leprae* compared to *M. tuberculosis* [36]; and ML0247, a putative arsenate reductase arsC. Using the template of reactions available for *M. tuberculosis* H37Rv in GSMN-TB_2 (856 reactions catalysed by 729 enzymes), we were able to transfer 713 reactions to construct the *M. leprae* model using gene orthology. However, only 396 of orthologues appeared to be functional in *M. leprae*, the remainders were pseudogenes. 21 additional reactions were added to the *M. leprae* model based on manual literature curation, and the resulting *M. leprae* network encoded a total of 417 genes encoding metabolic enzymes. This reduced the number of functional enzymes (nearly half the number as compared to *M. tuberculosis*), which is a direct consequence of the massive genome reduction as well as pseudogene formation in *M. leprae*.

### Incorporation of pseudogenes and orphan reactions facilitated *in silico* growth of GSMN-ML

We applied flux balance analysis (FBA) to investigate the ability of the *in silico M. leprae* model to synthesize biomass from basic nutrients, starting from the minimal nutrient requirements (supplementary File S2) needed to grow *M. tuberculosis in vitro*. GSMN-ML was however unable to synthesize biomass from the minimal requirements that are needed by GSMN-TB. To address this problem we reintroduced reactions that were removed in constructing GSMN-ML from GSMN-TB (because genes encoding these reactions were pseudogenes in *M. leprae*) until we obtained a feasible network. This resulted in the introduction of several orphan reactions that appeared to be essential in *M. leprae*, but whose function is not encoded by any known functional *M. leprae* gene. We then adopted two approaches to minimize the number of these GSMN-ML orphan reactions. First, we performed bioinformatic analysis to identify any hypothetical gene that could complement a function provided by an essential orphan reaction. If such a gene was found, it was added to the network. However, in many cases, no putative gene could be identified to potentially encode the function of the orphan gene. We then investigated whether the orphan reaction could be made non-essential by opening a transport gate for a metabolite that was potentially imported from the intracellular environment of *M. leprae*. If that was the case then we opened that transport gate to allow import and removed that orphan reaction from the network. Lastly, essential orphan reactions that could not be complemented by a hypothetical gene whose product could not be feasibly obtained from the host were left as functional orphan reactions. This operation led to the generation of a feasible network, GSMN-ML that included eight orphan transport reactions (Table 1). A full list of network reactions and the medium components used for this analysis is included in supplementary files S2 and S3 respectively.

**Table 1.** Medium supplements for growth of GSMN-ML. Transport gates for the metabolites shown are included in supplementary File S3.

| <u>Metabolite</u> | <u>Transport_gate</u> |
| --- | --- |
| L, L-diaminopimelate | R827 |
| Lysine | R834 |
| Methionine | R835 |
| Formate | R806 |
| Tetrahydrofolate | R936 |
| Pantothenate | R939 |
| COB-I | R937 |
| Protoporphyrinogen | R938 |

The following potentially imported nutrients that were identified by this procedure were formate, L,L-2,6-diaminopimelate, lysine, methionine, tetrahydrofolate, coenzyme-B (COB-I), pantothenate and protoporphyrinogen (Table 1). The requirement for methionine and lysine biosynthesis is caused by the genes *metC* encoding O-acetyl-L-homoserine sulfhydrylase (ML0683c) and *dapB* that encodes dihydrodipicolinate reductase (ML1527) being pseudogenes in *M. leprae*. The requirement for L,L-2,6-diaminopimelate (DAPIM), an intermediate for lysine synthesis and one of the principle substrates for peptidoglycan biosynthesis, is due to the absence of diaminopimelate decarboxylase (EC: 4.1.1.20) in *M. leprae* (Table 1). The requirement for tetrahydrofolate, formate and COB-I is due to many genes involved in their synthesis being pseudogenes in *M. leprae*. The requirement for pantothenate, the substrate for coenzyme-A (CoA) synthesis, is due to *M. leprae* lacking 2-dehydropantoate 2-reductase (EC: 1.1.1.169) that is needed to synthesize CoA from glycolysis. The requirement for protoporphyrinogen is a consequence of *M. leprae*’s inability to synthesise porphyrin, and thereby any hemoproteins, from L-glutamate due to the absence of the hemN gene, coding for oxygen-independent coproporphyrinogen III oxidase. The import flux of these nutrients was fixed to the minimum needed for maximum biomass production (see supplementary File S3 and Figure 1).

**Figure 1:**
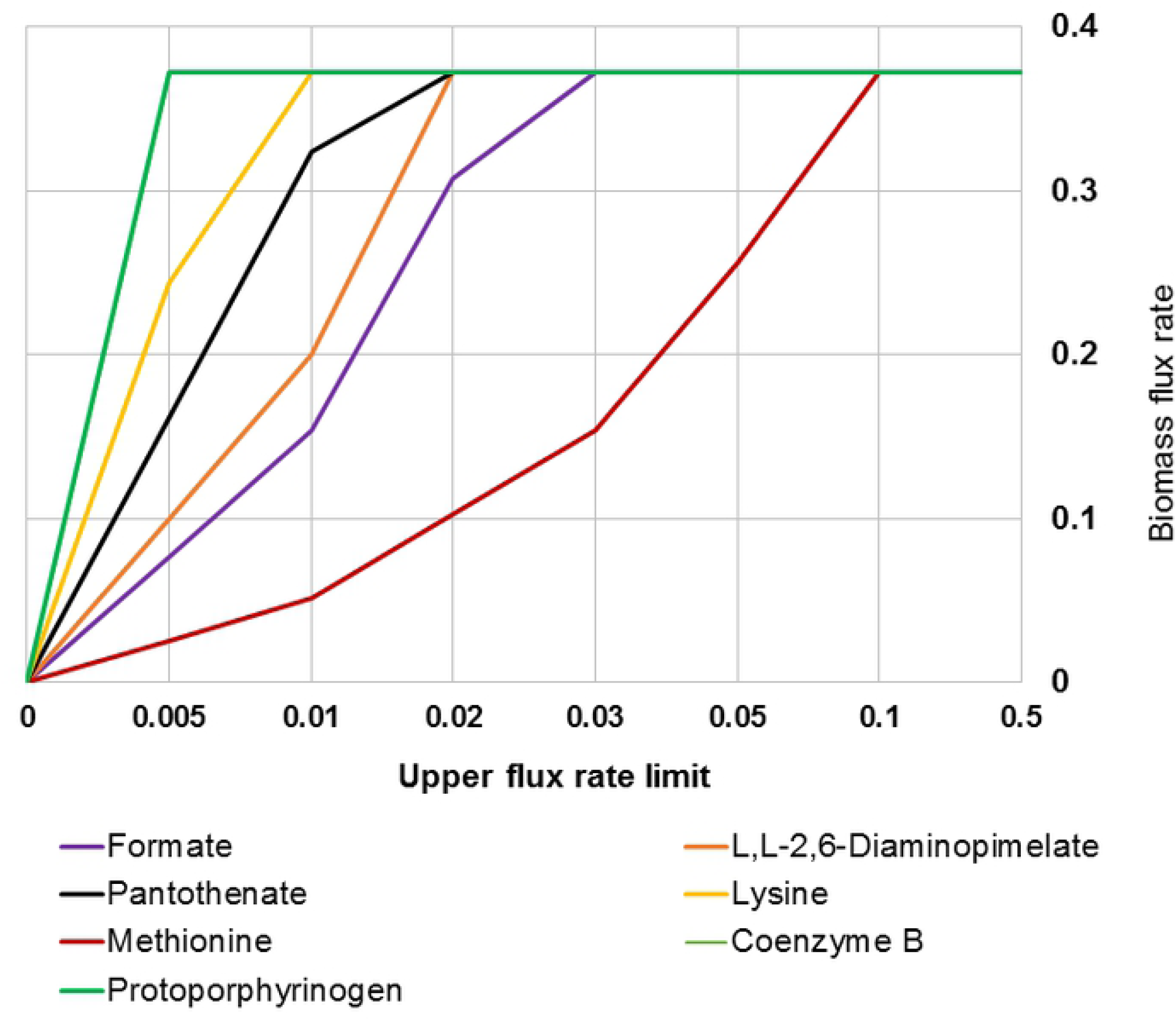
Influence of opening transport gates for essential *in silico* medium supplements on predicted biomass production for the *M. leprae* GSMN model. Media component uptake was limited to maximal value of 1 mmol g^-1^dryweight h^-1^. The biomass flux rate for GSMN-ML was calculated with respect to the media components

### Substrate utilization by GSMN-ML

A major finding of the genome sequencing project by Cole *et al*., 2001 was that *M. leprae* has lost many catabolic genes and pathways [10]. To investigate this in more detail we investigated the ability GSMN-ML to utilize various carbon and nitrogen substrates for biomass production and compared it to GSMN-TB_2 (supplementary File S4). The results, (Figure 2A) shows that, perhaps surprisingly, *M. leprae* has retained the ability to utilize many carbon sources; yet, in comparison with *M. tuberculosis*, has lost the ability to utilize acetate and glycolate due to pseudogenization of acetate kinase, phosphate acetate transferase and acetyl-coA synthase. *M. leprae* has also lost almost the entire pathway involved in cholesterol utilization and cannot thereby utilize host-derived cholesterol [37], which appears to be a growth substrate for *M. tuberculosis in vivo*. Catabolism of amino acids, such as valine, threonine and isoleucine as carbon sources was also impaired in *M. leprae* (Figure 2B). To explore the nitrogen requirements of GSMN-ML, FBA was performed with five different single nitrogen sources: ammonia, nitrate, urea, glutamate and glutamine (Figure 2C) and with glucose as the primary carbon source, comparing to GSMN-TB_2. Both models were able to utilize glutamate and glutamine to generate approximately equal biomass. The GSMN-TB_2 model was able to generate 31.3% more BIOMASS using ammonia as compared to GSMN-ML, and unlike *M. leprae*, was able to grow on urea as a single nitrogen source. Neither of the models could generate biomass using nitrate as nitrogen source. These data suggest that glutamate or glutamine is the most likely nitrogen source for *M. leprae*.

**Figure 2:**
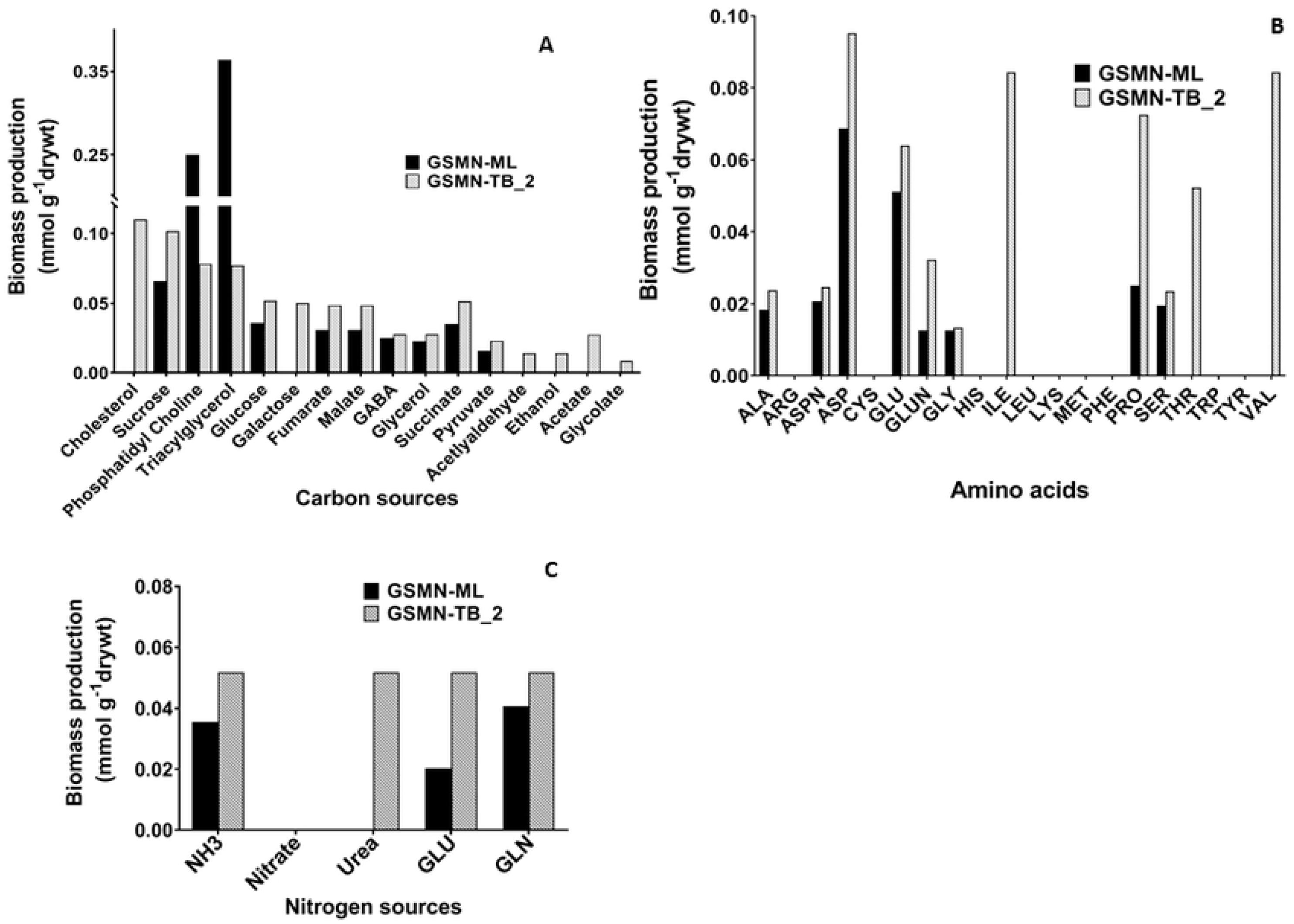
*In silico* prediction of biomass production for various substrates of *M. leprae* compared to the *M. tuberculosis* GSMN models. Carbon substrate utilization profiles of *M. leprae* GSMN-ML vs. *M. tuberculosis* GSMN-TB_2. Biomass production is shown for both the models on (A) various carbon substrates and (B) amino acids as carbon sources (C) various nitrogen sources. Biomass production flux was calculated on the various substrates using FBA. Abbreviations for amino acids are alanine (ALA), glycine (GLY), arginine (ARG), asparagine (ASPN) aspartic acid (ASP), cysterine (CYS), glutamic acid (GLU), glutamine (GLUN), histidine (HIS), isoleucine (ILE), leucine (LEU), lysine (LYS), methionine (MET), phenylalanine (PHE), proline (PRO), serine (SER), threonine (THR), tryptophan (TRP), tyrosine (TYR) and valine (VAL).

### DPA to predict intracellular metabolism of *M. leprae* from transcriptome data

Directly measuring metabolism is very difficult, particularly for intracellular bacteria. It is however relatively straightforward to obtain gene expression profiles of intracellular bacteria and, from that, discover whether key metabolic genes are up- or down-regulated in a certain condition. However, single genes may contribute to several different metabolic pathways; and most metabolic pathways involve many genes, which may show contradictory expression patterns. To provide a systems-wide insight into intracellular metabolism, we previously developed Differential Producibility Analysis (DPA) that uses a genome scale model to associate metabolites with system-wide analysis of gene expression patterns [20]. The first step in DPA is to identify all genes (the metabolite producibility gene list) that are required (in the genome-scale model) for production of every metabolite, given a set of nutrients inputs. The next step in DPA is to interrogate gene expression data to identify whether the genes that contribute to the production of a particular metabolite tend to be associated with either up-regulated or down-regulated genes in the test condition. Note that metabolites may appear in both lists if, for example, one set of genes associated with their production is found to be up-regulated and another set is found to be down-regulated. In our previous study, we applied DPA to gene expression data obtained from both *E. coli* and *M. tuberculosis*, including *M. tuberculosis* replicating within macrophages [20]. In this study, we applied the same approach to investigate metabolic changes associated with *in vivo* growth of *M. leprae*.

Since the nutrients used by *M. leprae in vivo* are unknown, we had to perform the first step of DPA with putative intracellular nutrients. Enzymes for catabolism of hexoses and glycerol have been detected in *M. leprae* isolated from armadillos [38, 39], prompting us to perform the DPA growth simulations with both (independently) glucose and glycerol. RNA was extracted and immediately processed from *in vivo*-grown viable *M. leprae* harvested and from mouse footpads [21] (see Methodology for details). The control condition was *M. leprae* from mouse footpads, as above, but then incubated in axenic culture for 48 hours under conditions (see Methodology) where the bacilli remain viable but do not replicate. *In vivo* up-regulated genes were defined as being the product of an enzyme transcribed from a gene with a fold change >2 in the RNA-Seq data. *In vivo* down-regulated genes are defined as genes with a fold change <0.5 in the RNA-Seq data (supplementary File S5). These expression lists were then cross-checked against the metabolite genes to identify metabolite whose production (genes encoding the enzymes involved in their production) is significantly associated with the up-regulated gene list, or is significantly associated with the down-regulated gene list. The top 100 metabolites associated with up- or down-regulated genes for both carbon sources is presented in supplementary File S6. For up-regulated genes, 67 and 54 metabolites were shared between GSMN-ML using glucose and glycerol as carbon sources (Figure 3A, B). Pathways and areas of metabolism likely to be affected by up and down-regulated genes in *M. leprae* during growth on the two carbon sources are illustrated in Figure 3C-F. Metabolites required for synthesis of amino acids comprised the major category of metabolites predominantly associated with up-regulated genes in both glucose and glycerol; whereas metabolites involved in cofactor synthesis were mostly associated with upregulated on glucose simulations (Figure 3C, D). Metabolites involved in lipid synthesis were the major family of metabolites identified as being associated with down-regulated genes for growth on glucose (Figure 3E). On glycerol, metabolites for co-factor biosynthesis were down-regulated on glycerol (Figure 3F). Several metabolites involved in cell wall synthesis of *M. leprae*’s cell wall were associated with up-regulated genes only during glycerol utilisation simulations. This is likely because cell wall components, such as arabinogalactan-decaprenyl phosphate [40], and d-alanyl-d-alanine, a peptide-glycan precursor [41], fatty-acid derived mycolates and arabinogalactan-peptidoglycan molecules are derived directly from glycerol metabolism.

**Figure 3:**
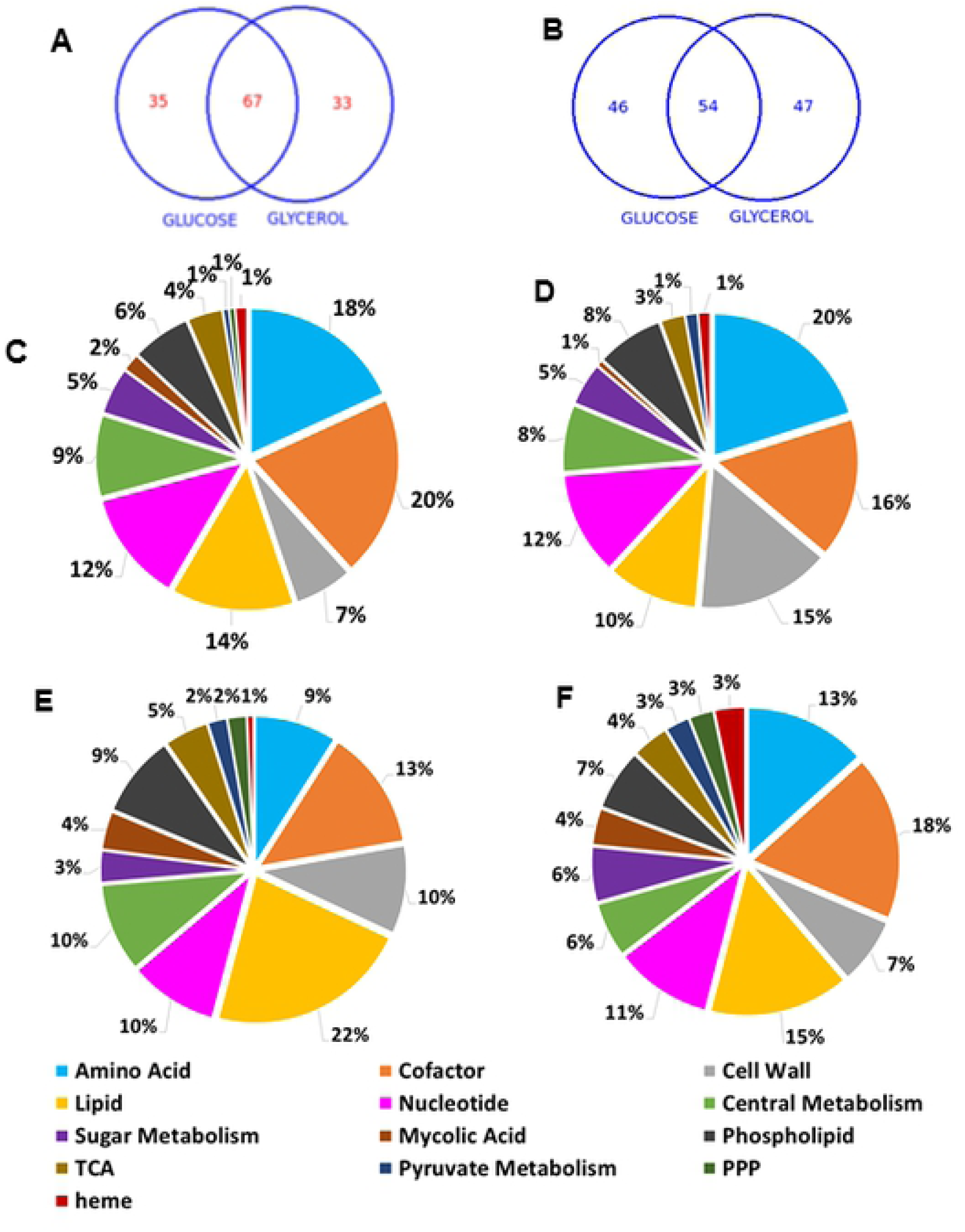
DPA analysis of gene expression patterns to identify metabolic processes associated with up-regulated (A) and down-regulated *M. leprae* genes (B) *in vivo*. Venn diagram compares metabolites identified as associated with up-regulated (C & D) or down-regulated (E and F) genes using either glucose (C & E) or glycerol (D & F) as carbon source in DPA.

This pattern of gene regulation of *M. leprae* in response to its *in vivo* environment appears to be very different from that of intracellular *M. tuberculosis* [20]. For example, in *M. tuberculosis* 25% of metabolites associated with phospholipid synthesis were associated with upregulated genes compared to only 1% associated with down-regulated genes [20]; whereas, in *M. leprae*, for glycerol and glucose-based simulations, a similar fraction of around 7% of metabolites involved in phospholipid synthesis were found to be associated with both up and down-regulated genes. Also, in *M. tuberculosis*, there was a clear tendency for metabolites involved in lipid metabolism (other than phospholipids) to be associated with up-regulated genes, whereas, for *M. leprae*, they were mostly associated with down-regulated genes. This pattern was reversed for amino acid metabolism, with a clear association with down-regulated genes in *M. tuberculosis* (25% compared with 15% associated with up-regulated genes) [20], whereas, in *M. leprae*, 18% of metabolites involved in amino acid synthesis were associated with up-regulated genes and only 9% associated with down-regulated genes. Comparing the two pathogens, the adaptation of *M. tuberculosis* to its intracellular environment appears to be associated with an up-regulation of lipid synthesis and down-regulation of amino acid synthesis; whereas the reverse appears to be true for *in vivo M. leprae*.

To assess the importance of *M. leprae* genes, i.e., whether a gene is essential or non-essential for growth on glycerol and glucose, we used GSMN-ML for gene knockout predictions. We calculated the growth rate when a gene in the GSMN-ML network was removed. We calculated gene essentiality (GS) ratios- the ratio of the growth rate of the knock out to that of wildtype for each gene in the network (supplementary Files S7, S8). Figure 4, 5 shows the central carbon network genes and the GS ratios. A gene was considered essential if the GS ratio was < 0.01. We found 21 genes involved in glycolysis, gluconeogenesis, TCA cycle and PPP to be essential central metabolic genes for growth of *M. leprae* on both glucose and glycerol. We compared the GS ratios for the central metabolic network genes with that of *M. tuberculosis*, by performing GS analysis of GSMN-TB_2 using glucose or glycerol as the sole carbon source. Around 64-66% genes of the total genes in the network were predicted to be essential in GSMN-ML compared to ∼28% in GSMN-TB_2. The higher percentage of essential metabolic genes in *M. leprae* is likely a consequence of the loss of non-essential genes due to deletion or pseudogenization during evolution of *M. leprae* from it progenitor.

**Figure 4:**
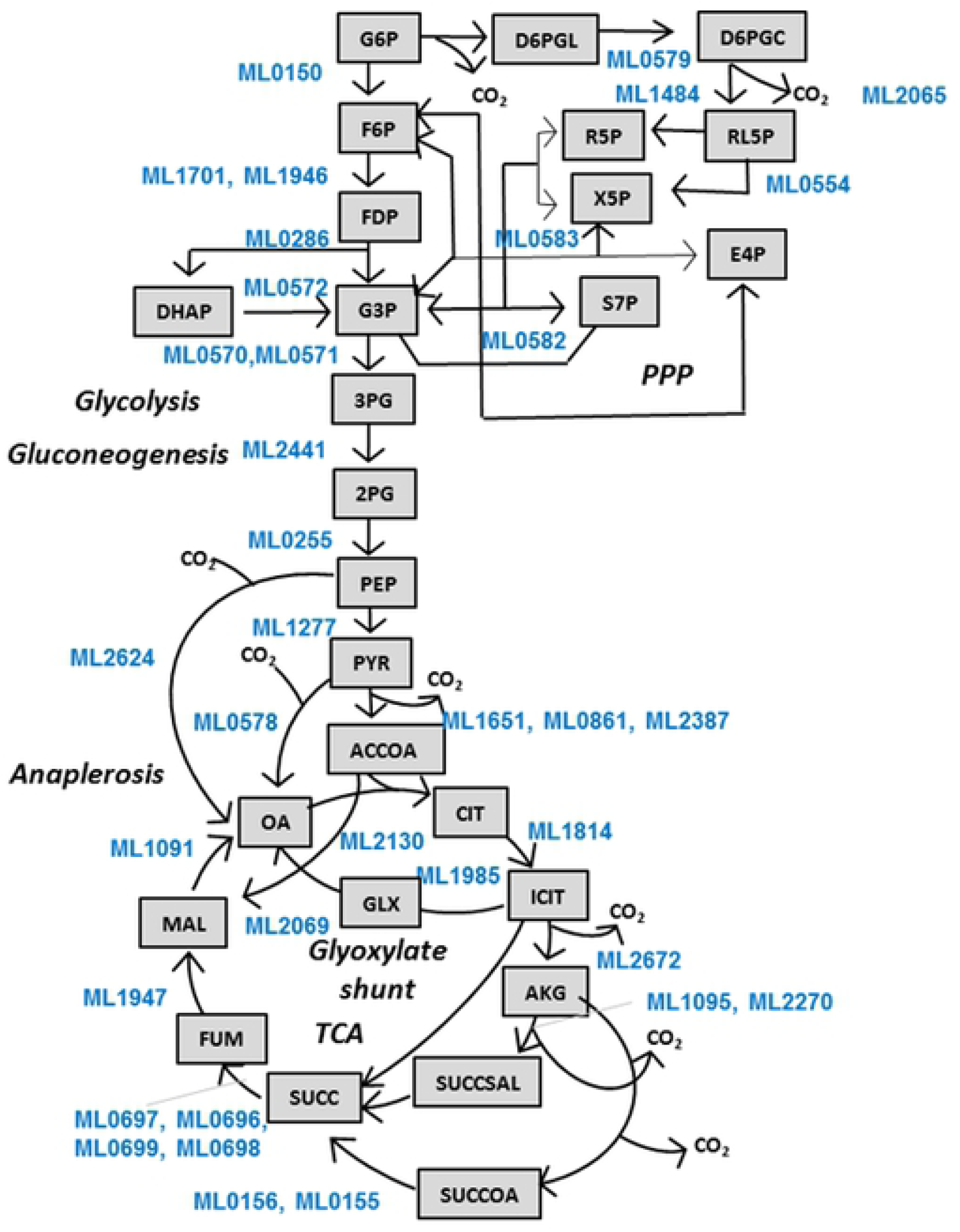
Central metabolic network of GSMN-ML. Reactions and their respective annotated genes (highlighted in blue) are shown for Glycolysis, Gluconeogenesis, PPP, TCA cycle, Glyoxylate shunt and anaplerotic CO2 fixation. Abbreviations for metabolites are: G6P (glucose-6-phosphate), F6P (fructose-6-phosphate), FDP (fructose-1,6-bisphosphate), G3P (D-glyceraldehyde-3-phosphate), DHAP (dihydroxyacetone-phosphate), 3PG (3-phosphoglycerate), 2PG (2-phosphoglycerate), PEP (phosphoenolpyruvate), PYR (pyruvate), ACCOA (acetyl-CoA), OA (oxaloacetate), MAL (malate), FUM (fumarate), SUCC (succinate), SUCCOA (succinyl-CoA), SUCCSAL (succinate_semialdehyde), AKG (α-ketoglutarate), ICIT (isocitrate), CIT (citrate), GLX (glyoxylate), D6PGL (D-gluconolactone-6-phosphate), D6PGC (6-phospho-D-gluconate), R5P (D-ribose-5-phosphate), RL5P (D-ribulose-5-phosphate), X5P (D-xylulose-5-phosphate), E4P (D-erythrose-4-phosphate) and S7P (D-sedoheptulose-7-phosphate).

**Figure 5:**
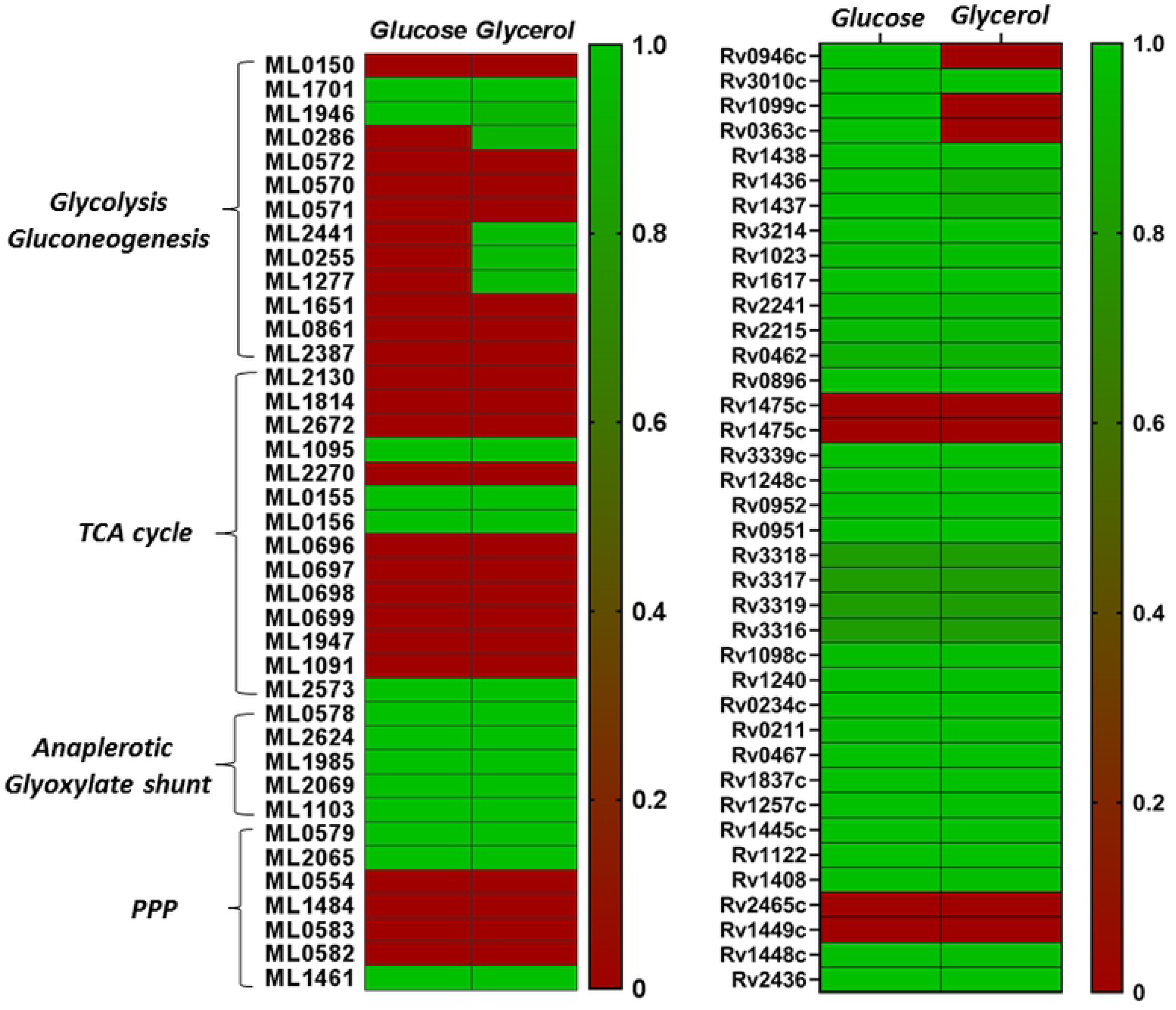
Gene essentiality analysis of *M. leprae* and *M. tuberculosis* utilizing glucose and glycerol as carbon sources. Heat map showing the gene essentiality (GS) predictions of central carbon metabolic genes during growth of *M. leprae* on glucose and glycerol. Simulations were performed in SurreyFBA platform and a gene was considered essential if the GS ratio < 0.01. Essentiality is indicated by single colour gradient with the maximum GS ratio of 1 (green) and minimum GS ratio (red).

## Discussion

In this study, we constructed GSMN-ML, a genome scale metabolic model of *M. leprae*. Interestingly, the model predicted that the pathogen would be unable to grow without the addition of supplements (Table 1) that the pathogen probably obtains from the host *in vivo*. Although we predict that these supplements are required for *in vitro* growth of *M. leprae*, it they are likely to be present in many of the media that have been tested in attempts to grow *M. leprae in vitro*. So it seems likely that additional, as yet unidentified, vital nutrients are needed for *in vitro* growth of the pathogen. Nonetheless, Table 1 does at least provide a minimal nutrient requirement that could be further investigated. Host-derived lipids are a well-established carbon source in mycobacterial infection such as *M. tuberculosis* [42-44]. Although there is no direct evidence of *M. leprae* using host cell lipids as a nutrient source, *M. leprae* infection did induce changes in lipid metabolism in the host Schwann cells and macrophages, making them lipid-rich foamy cells [45-47]. Also, lipid droplet biogenesis in the host provides growth advantage to *M. leprae* by providing lipid nutrient pools to the pathogen [46-48]. Our FBA-based simulations indicated that carbon sources triacylglycerol and phosphotidyl choline provided the highest biomass production for *in silico* growth on GSMN-ML, suggesting these, or similar lipids, as potential carbon sources for *M. leprae*. In comparison to *M. tuberculosis*, GSMN-ML failed to use either amino acids or a range of other lipid-derived metabolites as carbon sources. Similarly, and in contrast to *M. tuberculosis*, GSMN-ML was incapable of *in silico* growth with cholesterol as a carbon source (Figure 2A). This is in agreement with previous studies that indicate that, despite retaining an ability to oxidize cholesterol, *M. leprae* is unable to use cholesterol as an energy or carbon source [37]. This is consistent with the genome studies, which indicate that evolutionary gene-decay in *M. leprae* has resulted in the loss of the mce4 operon and that encodes for active transport system for cholesterol [10], [37], as well as many genes required for cholesterol catabolism. Despite being part of a major component to mycobacterial cell walls [48], GSMN-ML also failed to utilize galactose; indicating that galactose-cell wall components are likely synthesised by the Leloir pathway in the leprosy bacillus from UDP-glucose, rather than from galactose directly [10], [11]. Amino acid utilisation in *M. leprae* and *M. tuberculosis* were also significantly different. While *M. leprae* retains the ability to biosynthesize amino acids, catabolic pathways are degraded, accounting for why the pathogen cannot utilize amino acids as carbon sources (Figure 2B) [10]. This deficit is largely explained by the absence of the urease operon in *leprae*; also accounting for the inability of GSMN-ML to utilize urea as the nitrogen source (Figure 2C).

Our FBA analysis of GSMN-ML indicates that *M. leprae* is able to utilize both glucose and glycerol as carbon sources, consistent with evidence that glucose and/or glucose-derived metabolites were used as nutrients by *M. leprae* during *in vivo* growth [38], [49]. The finding may also be relevant to the observation that glucose metabolism of host Schwann cells is perturbed by *M. leprae* infection [49]. DPA analysis of gene expression data identified clear differences in intracellular metabolism of *M. leprae* compared to intracellular *M. tuberculosis*, particularly in lipid and amino acid metabolism. These differences may contribute to the very different virulence properties of these two related pathogens. Lastly, our analysis confirmed the expected higher degree of gene essentiality in *M. leprae* compared to *M. tuberculosis*. This is likely to be associated with a lower degree of metabolic robustness in the leprosy pathogen, compared to the TB pathogen; a vulnerability that might provide novel opportunities for new drugs.

## Conclusion

Despite intensive efforts, *M. leprae* has never been grown in the laboratory, severely hampering research on this important pathogen. In this study, we describe the construction of an *in silico* genome scale metabolic model of *M. leprae* GSMN-ML that is able to simulate growth. The model was used to devise a base medium for efforts to grow the pathogen in the laboratory. We also used the model to interrogate gene expression data from *in vivo*-grown *M. leprae* and identified several metabolic differences between that and *in vitro*-incubated cells. We also found evidence for major differences in the intracellular metabolism of *M. leprae* compared to the related pathogen, *M. tuberculosis*. The availability of the GSMN-ML model provides a new tool for the further exploration of the biology of this important pathogen that could lead to development of new approaches for control of leprosy.

## Acknowledgement

We thank BBSRC grant: BB/L022869/1, MRC/Newton Fund: MR/M026434/1, BBSRC: India Partnering Award IPA1825 and Bioinformatics Resources and Applications Facility (BRAF) of Government of India for funding. We thank National Institute of Allergy and Infectious Diseases (NIAID) Interagency Agreement IAA 15006-004-00000 for the provision of viable *M. leprae* for this study.

## Main Table

Table 1: **Essential *in silico* medium supplements and their transport gates in GSMN-ML**.

## Supplementary information

Supplementary File S1. **Metabolic network of GSMN-TB_2**. The network file includes metabolite lists, pathways and reactions for *M. tuberculosis* and the problem file (p-file) including the list of medium supplements and the upper and lower limits of each medium component uptake used for nutrient utilization analysis. The network was constructed using SurreyFBA software.

Supplementary File S2. ***In silico* medium supplement composition**. This composition was used to test and incorporate pseudogenes and orphan reactions that facilitated the growth of GSMN-ML. The p-file includes the list of medium supplements and the upper and lower limits of each medium component uptake used for nutrient utilization analysis.

Supplementary File S3. **Metabolic network of GSMN-ML**. The network file includes metabolite lists, pathways and reactions for *M. leprae* and the problem file (p-file) including the list of medium supplements and the upper and lower limits of each medium component uptake used for nutrient utilization analysis. The network was constructed using SurreyFBA software.

Supplementary File S4. **P-files used in SurreyFBA to generate growth and biomass production with GSMN-ML and GSMN-TB_2 model.** The p-files include the list of nutrients and their uptake rates-upper and lower limits.

Supplementary File S5. **RNA-seq data for *M. leprae* isolated from mouse foot pads.** The file includes expression data and fold changes in expression of M. leprae freshly harvested vs. 48h incubation in axenic medium.

Supplementary File S6. **List of up-regulated and down-regulated metabolites identified using DPA analysis.** Metabolites are listed for GSMN-ML and RNA-seq analysis using glucose and glycerol as sole carbon sources. The p-files include the list of nutrients and their uptake rates-upper and lower limits.

Supplementary File S7. **Flux balance analysis of GSMN-ML on glucose and glycerol as carbon sources and gene essentiality predictions.** File shows flux variability analysis and gene essentiality analysis for the model. Two media p-files was set for this test. One is to use glycerol as only carbon source and the other is to use glycerol as only carbon source and the other necessary molecules. Table ‘FVA’ shows results for all reactions with maximised biomass. Table ‘media’ shows media settings, those substrates whose lower-bound < 0 are allowed to be uptaken. Table ‘metabolite_name’ shows metabolite IDs and their names. Table ‘KO_gene’ shows gene essentiality analysis results on the media set in ‘Media’. GRateKO: growth rate when the gene knocked out. Ratio: calculated as GRateKO/GRateWT(wildType). Genes are considered as essential If Ratio < 0.01 (marked red).

Supplementary File S8. **Flux balance analysis of GSMN-TB_2 on glucose and glycerol as carbon sources and gene essentiality predictions.** File shows flux variability analysis and gene essentiality analysis for the model. Two media p-files was set for this test. One is to use glycerol as only carbon source and the other is to use glycerol as only carbon source and the other necessary molecules. Table ‘FVA’ shows results for all reactions with maximised biomass. Table ‘media’ shows media settings, those substrates whose lower-bound < 0 are allowed to be uptaken. Table ‘metabolite_name’ shows metabolite IDs and their names. Table ‘KO_gene’ shows gene essentiality analysis results on the media set in ‘Media’. GRateKO: growth rate when the gene knocked out. Ratio: calculated as GRateKO/GRateWT(wildType). Genes are considered as essential If Ratio < 0.01 (marked red).

